# svtools: population-scale analysis of structural variation

**DOI:** 10.1101/494203

**Authors:** David E. Larson, Haley J. Abel, Colby Chiang, Abhijit Badve, Indraniel Das, James M. Eldred, Ryan M. Layer, Ira M. Hall

## Abstract

**Summary:** Large-scale human genetics studies are now employing whole genome sequencing with the goal of conducting comprehensive trait mapping analyses of all forms of genome variation. However, methods for structural variation (SV) analysis have lagged far behind those for smaller scale variants, and there is an urgent need to develop more efficient tools that scale to the size of human populations. Here, we present a fast and highly scalable software toolkit (svtools) and cloud-based pipeline for assembling high quality SV maps – including deletions, duplications, mobile element insertions, inversions, and other rearrangements – in many thousands of human genomes. We show that this pipeline achieves similar variant detection performance to established per-sample methods (e.g., via LUMPY), while providing fast and affordable joint analysis at the scale of ≥100,000 genomes. These tools will help enable the next generation of human genetics studies.

**Availability and Implementation:** svtools is implemented in Python and freely available (MIT) from https://github.com/hall-lab/svtools.

## INTRODUCTION

With the dramatic cost reduction of whole genome sequencing (WGS) in recent years, large-scale human genetics studies are underway that aim to conduct comprehensive trait association analyses in tens to hundreds of thousands of deeply sequenced (>20x) individuals. Foremost among these are NIH programs such as NHGRI’s Centers for Common Disease Genomics (CCDG) and NHLBI’s Trans-Omics for Precision Medicine (TOPMed), which have generated >150,000 deep WGS datasets thus far. Moreover, ongoing genome aggregation efforts seek to produce even larger genome variation maps that can be mined for insights into genome biology, and to help interpret personal genomes and rare disease studies. These efforts, along with many others around the world, will usher in a new era of data-centric human genetics research.

A key promise of WGS is the potential to assess all forms of genome variation. However, despite considerable effort and creativity by many groups (most notably the 1000 Genomes Project, 1KGP (Mills, et al., 2011; Sudmant, et al., 2015)), it remains extremely difficult to assemble high quality SV maps from WGS data, especially for large cohorts comprising thousands of individuals. An initial obstacle is that SV detection is a difficult problem even for small-scale studies due to fundamental limitations in the integrity of short-read alignment signals used to infer breakpoint positions and estimate copy number from Illumina WGS data. These alignment signals – including split-read (SR), clipped-read (CR), read-pair (RP), and read-depth (RD) – are difficult to distinguish from sequencing and alignment artifacts, and are difficult to integrate with each other, such that even the best performing tools generally suffer from low sensitivity, high false discovery rates (FDR), and high compute costs.

A second issue is that current WGS-based multi-sample variant detection approaches require “joint” analysis of raw (or nearly raw) alignment data for each sample, at each putative variant site; however, due to memory and compute limitations, native joint calling algorithms do not perform well beyond the scale of several hundred genomes. SNV/indel detection tools such as GATK and VT have implemented distributed workflows to distill and combine variant detection signals in large cohorts through the use of intermediate files (e.g., gVCF) and parallel “scatter-gather” computing schemes. A natural goal is to develop similar approaches for SVs.

However, for SV the problem is different and arguably much harder. Tools must tolerate higher error rates and accommodate diverse variant sizes and architectures including balanced, complex and/or repetitive variants that may be difficult to classify. Parallelization schemes are complicated by the fact that, unlike SNVs that map to a *single* coordinate, SV breakpoints are defined by *pairs* of discontiguous and potentially distant strand-oriented reference genome coordinates (formally “break-ends”) that describe novel DNA junctions in the experimental genome (formally “novel adjacencies”). Intermediate data structures analogous to gVCF are difficult to design because they must encapsulate information from at least four disparate alignment signals (SR, CR, RP, RD), each with different resolution and artifact modalities. Cross-sample merging schemes must be robust to positional uncertainty because SV breakpoint mapping resolution is typically imprecise on a per-sample basis (~10-100 bp mean), and sequencing and alignment effects can vary widely across variant classes, samples and batches. New approaches are required.

Of course, the task of combining spatially imprecise SV/CNV calls across collections of samples is an old problem that has been dealt with effectively through ad-hoc methods in prior microarray (Conrad, et al., 2009; Redon, et al., 2006; Wellcome Trust Case Control, et al., 2010) and WGS-based (Mills, et al., 2011; Sudmant, et al., 2015) studies. However, array-based methods do not readily extend to balanced SVs or to the increased resolution, complexity and scale of deep WGS. 1KGP employed a clever approach to merge results from multiple algorithms and platforms (Mills, et al., 2011; Sudmant, et al., 2015), but this was a monumental effort and the methods therein are impractical for routine use. GenomeSTRiP has two published workflows for detecting SV in populations of samples, but both have limitations: an early version focuses on deletions and serially combines RP-based detection with RD genotyping (Handsaker, et al., 2011); a second RD-based CNV pipeline is computationally expensive, low resolution (>1 kb), and limited to moderate sample sizes (<1,000) (Handsaker, et al., 2015). To our knowledge, no publicly available tools or reproducible workflows exist to systematically assemble high-resolution SV callsets from joint analysis of multiple alignment signals in tens of thousands of deep WGS datasets, as we present here.

## METHODS

We developed a software toolkit and distributed workflow for large-scale SV callset generation that combines per-sample variant discovery, resolution-aware cross-sample merging, breakpoint genotyping, copy number annotation, variant classification, and callset refinement (**Fig. 1**). We release the svtools python toolkit (https://github.com/hall-lab/svtools) and two pipeline versions: an on-premise “B37” version designed to handle BAM files aligned to the GRCh37 reference genome, that relies mainly on BASH scripts and LSF commands; and a cloud-based “B38” pipeline written in WDL designed to work with CRAM (Hsi-Yang Fritz, et al., 2011) files aligned to GRCh38 using the new “functional equivalence” standard developed by the CCDG and TOPMed programs (Regier, et al., 2018). Despite their different workflow implementations and reference genome versions, the core tools and parameters are virtually identical between these two pipelines, and both are publicly available (https://github.com/hall-lab/sv-pipeline).

**Figure 1.**
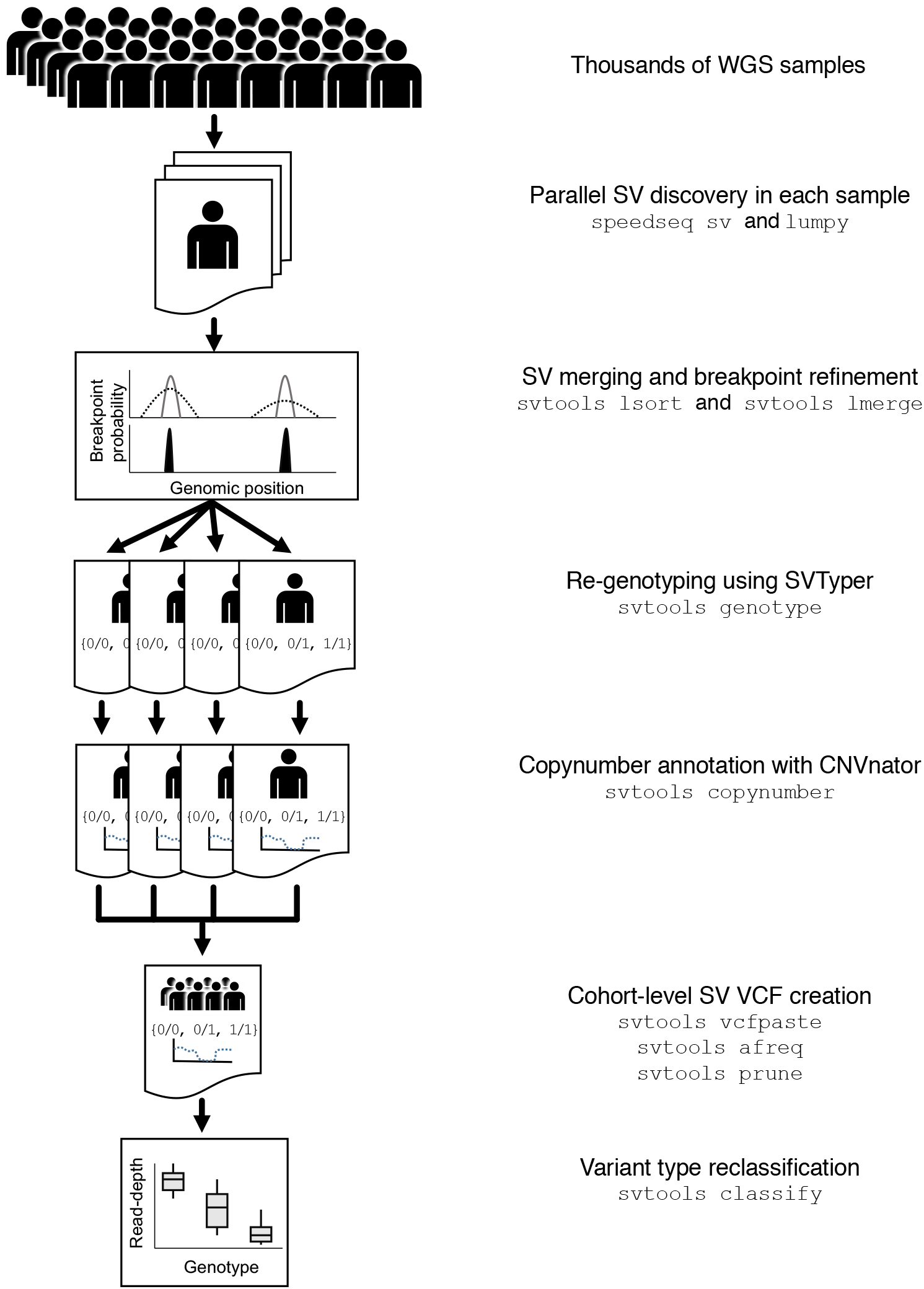
The svtools pipeline. SVs are detected separately in each sample using LUMPY. Breakpoint proPaPility distriPutions are utilized to merge and refine the coordinates of SV Preakpoints within a cohort, followed Py parallelized re-genotyping and copy numPer annotation. Variants are merged into a single cohort-level VCF file and variant types are classified using the compined Preakpoint genotype and read-depth information.

The first step is to analyze each genome separately, in parallel. We generate per-sample breakpoints calls using the LUMPY algorithm (Layer, et al., 2014), which combines RP and SR alignment signals in a probabilistic breakpoint detection framework. LUMPY is a widely used tool that has been benchmarked extensively in prior studies (Chiang, et al., 2015; Chiang, et al., 2017; Layer, et al., 2014); here, we adapted LUMPY to CRAM and improved performance on GRCh38 by masking highly repetitive and misassembled genomic regions. The second step is to merge all candidate variants from all samples into a single non-redundant VCF/BEDPE (Danecek, et al., 2011; Quinlan and Hall, 2010) file. Positional uncertainty is modeled during the merging process through the use of breakpoint probability distributions (as within LUMPY itself (Layer, et al., 2014)), which is possible because we have modified LUMPY to report the integrated per-base probability distribution for each breakpoint-containing confidence interval in the output per-sample VCF. The merging algorithm defines collections of SV predictions with mutually consistent coordinate intervals and orientations, as defined by the extent of overlap between breakpoint probability distributions in each sample, then combines and refines coordinates based on the weight of alignment evidence at each base, in each sample.

We then genotype all candidate SV breakpoints in all samples using SVTyper (Chiang, et al., 2015). This tool measures RP, SR and CR alignment signals at predicted breakpoints, in a more sensitive and accurate manner than feasible with genome-wide SV discovery tools such as LUMPY. The new svtools implementation handles CRAM and is significantly faster and more sensitive than the original (Chiang, et al., 2015). Quantitative allele balance information is retained throughout the workflow to preserve trait mapping power at difficult-to-genotype variants. Since certain CNVs are easier to genotype by RD analysis than breakpoint-spanning alignments, we also use CNVnator’s “genotype” tool (Abyzov, et al., 2011) to estimate the copy number of each SV interval in each sample. This copy number information can be used in lieu of breakpoint genotypes as desired in downstream association analyses, and is crucial for SV classification.

We next use a combination of breakpoint coordinates, breakpoint genotypes, read-depth evidence and genome annotations to classify each SV breakpoint either as a deletion (DEL), duplication (DUP), inversion (INV), mobile element insertion (MEI) or generic rearrangement of unknown architecture (“break-end”, BND) (see **Supplementary Methods** for details). This is an important and challenging step. The main difficulty arises because breakpoint prediction tools such as LUMPY are designed to detect novel DNA adjacencies, however, it is impossible to infer SV architecture from such evidence alone. For example, DEL and MEI variants often have identical breakpoint configurations (i.e., direct orientation), and complex rearrangements are often defined by multiple adjacent breakpoints that masquerade as simple SVs. The ‘svtools classify’ tool distinguishes DELs and DUPs from balanced SVs (BNDs) based on linear regression of quantitative breakpoint genotype information and copy number estimates from the affected genomic interval. MEIs are discerned by the location of mobile elements in the reference genome, and inversions by the co-detection of two breakpoints with inverted orientation. This classification step enables more informed SV impact prediction (Ganel, et al., 2017) and thus improves prioritization of rare SVs for human genetics studies. We note that additional SV classes – including interspersed duplications, retrogene insertions, translocations and complex rearrangements – can be identified by further interrogation of BND calls, but that rigorous automation of this process will require further work.

The final steps are callset refinement, tuning and quality control (QC). These are extremely important for obtaining high quality results, and ideally should take place with knowledge of genealogical relationships and SNVs. Miscellaneous tools are provided for allele frequency annotation, genotype refinement, confidence scoring, cross-callset variant look-ups, variant pruning, and file format conversion (see https://github.com/hall-lab/svtools#usage).

Additional details are available in the **Supplementary Methods**.

## RESULTS AND DISCUSSION

The sensitivity, accuracy and utility of the core SV discovery and genotyping algorithms in our pipeline have been thoroughly documented in multiple prior small-scale studies (Chiang, et al., 2015; Chiang, et al., 2017; Layer, et al., 2014). Here, we focus on the question of whether we achieve similar performance using the distributed workflow on much larger sample sizes. We constructed two separate callsets using identical methods: a “small” 12-sample callset composed solely of 1KGP samples, and a “large” 1000- sample callset composed of the same 1KGP samples plus 988 Finnish samples.

We first assessed the relative sensitivity obtained in the small vs. large callset using 1KGP calls as ground truth (**Table 1**). We achieved nearly identical sensitivity levels in per-sample calls prior to and after merging, at levels that are consistent with prior single-sample tests (Chiang, et al., 2015; Layer, et al., 2014). This demonstrates the effectiveness of our cross-sample merging strategy. Notably, sensitivity in both the small and large callsets improves markedly after the re-genotyping step. This demonstrates the benefits of re-genotyping, which is designed to be more sensitive than the initial SV discovery step and – when combined with high resolution cross-sample merging – emulates joint analysis by allowing evidence to be borrowed across samples. This important feature also provides quality and genotyping information for every sample, enabling confidence filtering of the variants. Taken together, this results in the uniform SV levels apparent in a 8,438-sample callset (generated for a different study) after re-genotyping and quality filtering, as compared to directly after calling (**Supplementary Fig. 1**). Sensitivity levels after re-genotyping are similar (if not better) in the large vs. small callset, which shows that our tools achieve comparable sensitivity at vastly different sample sizes. Note that these comparisons underestimate sensitivity given known false positives in 1KGP (Chiang, et al., 2015; Chiang, et al., 2017; Layer, et al., 2014).

**Table 1.** Detection sensitivity in large and small cohorts. Sensitivity is defined as percent of detectable 1000 Genomes Project variants identified in the cohort. HC stands for high confidence variants.

| Sample | 12 Sample Callset |  |  |
| --- | --- | --- | --- |
|  | Merge only | Reclassified (naïve bayes) |  |
|  | Sensitivity (all) | Sensitivity (all) | Sensitivity (HC) |
| HG00513 | 80.94% | 87.53% | 82.43% |
| HG00731 | 78.17% | 83.96% | 78.88% |
| HG00732 | 82.40% | 87.43% | 81.39% |
| NA12878 | 82.39% | 88.19% | 83.15% |
| NA19238 | 84.39% | 88.58% | 82.41% |
| NA19239 | 74.39% | 77.60% | 73.36% |

| Sample | 1000 Sample Callset |  |  |
| --- | --- | --- | --- |
|  | Merge only | Reclassified (regression) |  |
|  | Sensitivity (all) | Sensitivity (all) | Sensitivity (HC) |
| HG00513 | 80.23% | 88.03% | 83.80% |
| HG00731 | 77.67% | 84.47% | 80.46% |
| HG00732 | 81.50% | 88.12% | 82.56% |
| NA12878 | 81.58% | 88.62% | 84.18% |
| NA19238 | 83.86% | 88.53% | 82.80% |
| NA19239 | 74.01% | 77.81% | 73.31% |

We next examined variant calling accuracy. It is impossible to measure FDR in the absence of a comprehensive truth-set; however, Mendelian error (ME) rates are an informative proxy. To estimate ME, we examined inheritance patterns in 4 separate parent-offspring trios included in the 12-sample and 1000-sample callsets (**Table 2**). ME rates are high (12-17%) prior to re-genotyping, classification and confidence scoring, but fall to acceptable levels (2-3%) for high-confidence calls in the final small and large callsets. The slightly higher ME rate in the large callset is accompanied by substantially more variant calls and thus can be tuned to the desired ME rate depending of the application-specific desired balance between sensitivity and specificity.

**Table 2.** Mendelian error rate in large and small cohorts. Mendelian error (ME) rate is defined as the number of Mendelian errors divided by the total number of informative variants on the autosomes.

| 12 sample |  |  |  |  |  |  |
| --- | --- | --- | --- | --- | --- | --- |
| Merge only |  |  | Reclassified (naïve bayes) |  |  |  |
| All |  |  | All |  | High Confidence |  |
| Family | Variants | ME Rate % | Variants | ME Rate % | Variants | ME Rate % |
| CEPH1463 | 6107 | 12.72% | 6237 | 8.11% | 3184 | 2.29% |
| PR05 | 5783 | 15.75% | 6182 | 8.57% | 3164 | 2.24% |
| SH032 | 5670 | 16.83% | 6054 | 8.77% | 3182 | 2.64% |
| Y117 | 7534 | 15.18% | 7519 | 8.83% | 3889 | 2.24% |

| 1000 sample |  |  |  |  |  |  |
| --- | --- | --- | --- | --- | --- | --- |
| Merge only |  |  | Reclassified (regression) |  |  |  |
| All |  |  | All |  | High Confidence |  |
| Family | Variants | ME Rate % | Variants | ME Rate % | Variants | ME Rate % |
| CEPH1463 | 6147 | 12.93% | 10429 | 13.13% | 3605 | 2.77% |
| PR05 | 5827 | 15.99% | 10381 | 14.54% | 3629 | 2.98% |
| SH032 | 5708 | 16.92% | 10123 | 14.29% | 3574 | 3.33% |
| Y117 | 7568 | 15.38% | 11488 | 13.34% | 4208 | 2.69% |

Consistent with these results, the significantly larger callsets we have generated for other studies – based on 8,417 samples (on premises B37 pipeline) and 23,559 samples (cloud-based B38 pipeline) – exhibit similar numbers and types of variants (**Supplementary Fig. 1**) and achieve similarly low ME rates (data not shown). Taken together, these analyses demonstrate that our pipeline achieves high performance at large sample sizes.

A key strength of our pipeline is scalability and cost. Tool performance metrics are provided from sets of 10, 100, and 1,000 deep (>20x) genomes in **Table 3**. Overall, most steps are efficient and require modest compute resources, allowing them to be run on affordable cloud instances. For the 1,000 sample dataset used above, we estimate costs to be merely ~$0.30 per dataset. The initial per-sample SV discovery steps have been optimized for speed and cost (~$0.13 per genome) and scale linearly with the number of samples. Merging is a complex and compute-intensive process that can require significant RAM usage but is only necessary once per callset and can be parallelized. The current merging strategy is effective with as many as ~7,000 samples, using commodity hardware; however, for callsets exceeding several thousand samples we recommend a tiered scheme, whereby separate batches of data (e.g., ~1,000 samples) are combined during an initial sample-level merging step, followed by batch-level merging. Initial evidence suggests this approach results in similar if not higher quality breakpoint predictions than bulk merging (**Supplementary Fig. 2**), especially if data is batched by cohort and sequencing protocol.

**Table 3.** Computational benchmarking of svtools subcommands. For three different size cohorts, each tool was run (n=4) to generate mean CPU and RAM utilization. For the genotype and copynumber commands, benchmarking was performed on a single, representative sample within the cohort of median file size. All other commands were evaluated on the entire dataset. Some benchmarking runs finished before LSF was able to gather memory usage metrics and these are reported as NA.

| **Num. Samples** | **10** |  | **100** |  | **1000** |  |
| --- | --- | --- | --- | --- | --- | --- |
| **Program** | **CPU (m)** | **RAM (MB)** | **CPU (m)** | **RAM (MB)** | **CPU (m)** | **RAM (MB)** |
| **lsort** | 0.126 | 5.964 | 1.070 | 1696.008 | 11.610 | 3402.480 |
| **lmerge** | 2.113 | 87.791 | 18.752 | 258.402 | 194.322 | 2032.114 |
| **genotype** | 13.459 | 2008.828 | 31.774 | 1222.536 | 61.579 | 1255.593 |
| **copynumber** | 0.215 | NA | 0.285 | NA | 0.417 | NA |
| **vcfpaste** | 0.079 | NA | 1.382 | 75.660 | 68.035 | 181.845 |
| **afreq** | 0.098 | NA | 0.911 | 77.701 | 20.247 | 97.713 |
| **vcftobedpe** | 0.082 | NA | 0.138 | 3.277 | 0.824 | 70.851 |
| **bedpesort** | 0.082 | 17.363 | 0.182 | NA | 0.823 | 70.904 |
| **prune** | 0.175 | 18.790 | 0.402 | 61.667 | 1.365 | 171.728 |
| **bedpetovcf** | 0.081 | 17.368 | 0.177 | 34.794 | 0.756 | 71.003 |
| **vcfsort** | 0.038 | NA | 0.082 | 0.594 | 1.367 | 900.887 |
| **classify** | 6.983 | 530.722 | 8.278 | 526.480 | 25.806 | 680.900 |

The distributed genotyping step is the key bottleneck for large studies, since sensitive SV genotyping requires computationally expensive interrogation of raw alignment data, and aggregate compute time scales as a function of both sample size and the number of candidate variants. The latter is determined by a combination of sample size, ancestry composition, genetic relatedness, and per-sample variant discovery FDR, and is difficult to predict. Empirically, genotyping accounts for 6%, 12% and 24% of compute at the scale of 10, 100 and 1,000 genomes, respectively (**Table 3**), and ~78% of compute at the scale of 23,559 genomes.

Remarkably, a 23,559 genome callset was assembled on the Google Cloud at an empirical cost of $0.68 per sample. Based on the observed performance at different scales, we expect our current pipeline to achieve affordable callset generation (<$2 per sample) at the scale of ~10^5^ genomes, although improved methods may be necessary beyond that.

The tools described here will enable efficient and affordable analyses of SV in population-scale WGS studies, furthering our understanding of SV biology and enabling a more complete understanding of the contribution of SV to human traits.

## Supporting information

## ACKNOWLEDGEMENTS

We thank Aaron Quinlan and Brent Pedersen for discussion and suggestions; Allison Regier for critical contributions to SpeedSeq in support of svtools; and Krishna Kanchi for rigorous stress testing of the pipeline. We would also like to thank the users of the svtools package for bug reports, questions and suggestions that have greatly improved the software over time.

