## Supplementary material for "svtools: population-scale analysis of structural variation"

### Supplementary Figure 1

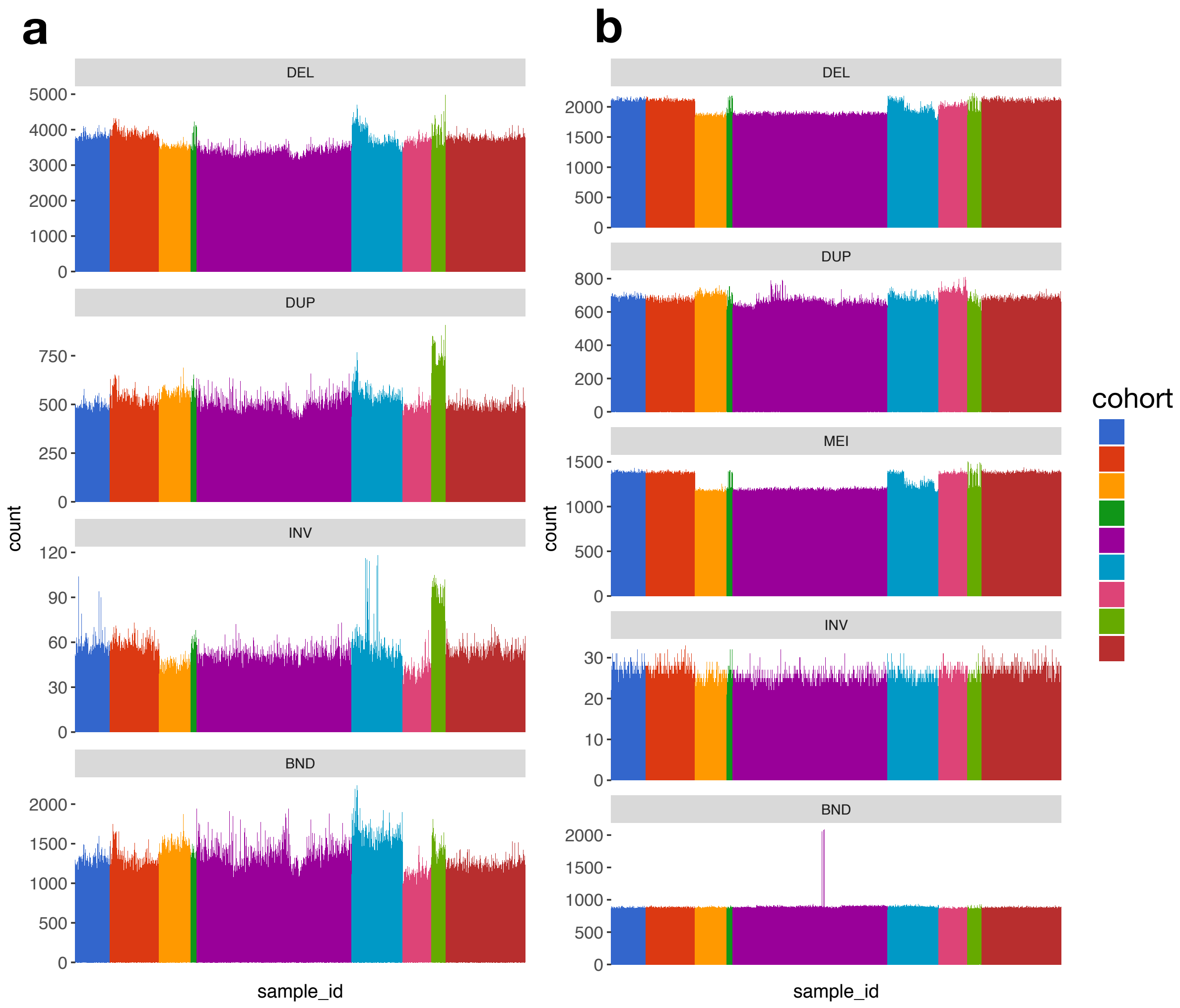


**Supplementary Figure 1.** Per-sample SV counts before and after merging. (a) Per-sample SV counts following the initial run of speedseq sv (restricted to autosomes only) in 8,438 samples. (b) High-quality SV counts per sample after completion of the svtools pipeline.

### Supplementary Figure 2

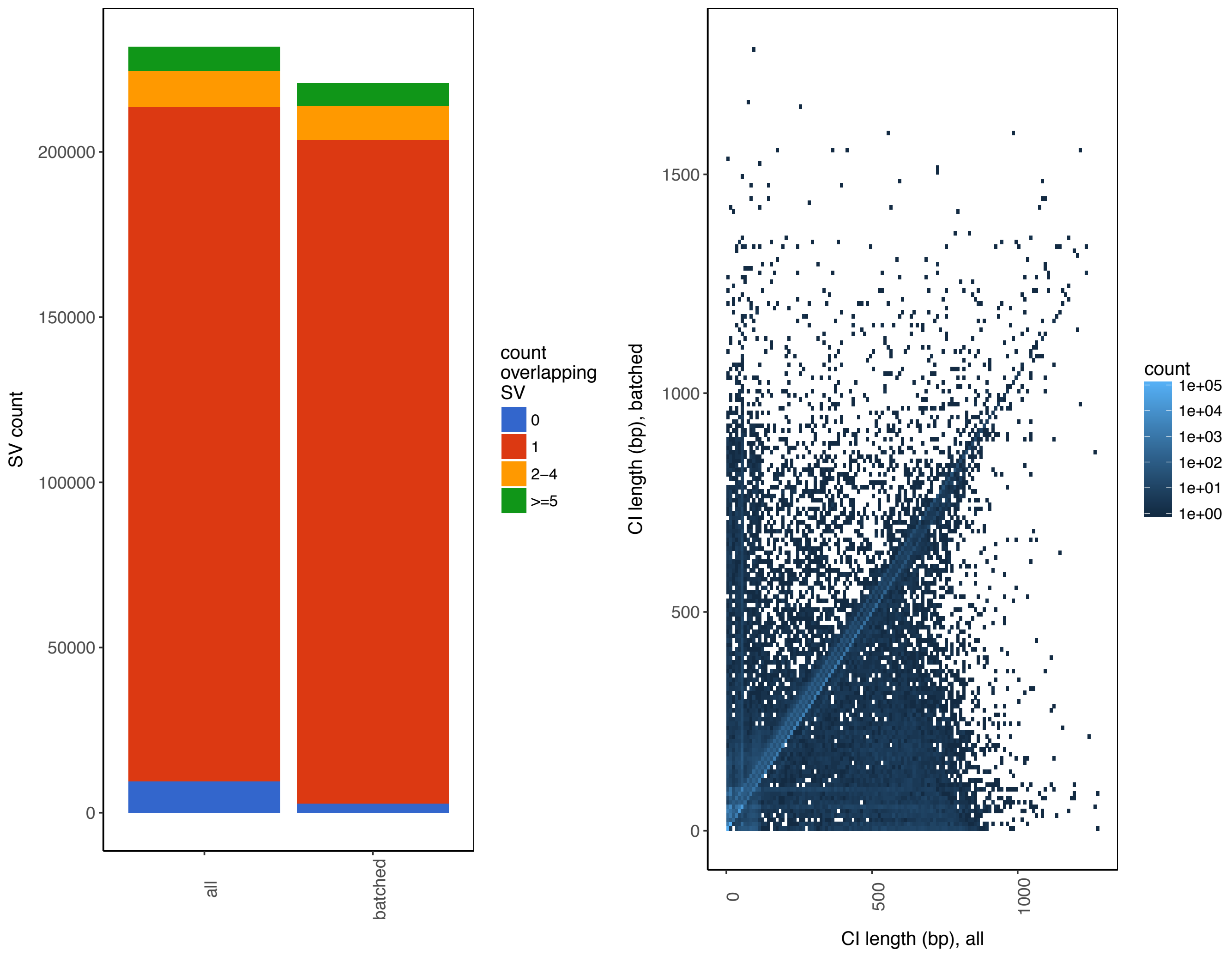


**Supplementary Figure 2.** Comparison of tiered and simple merging strategies. For both panels, 'All' indicates that all 8,438 samples were merged together in a single step and 'Batched' indicates that each of the nine cohorts was merged separately, followed by a second merged of the resultant files. (a) Overlap of SV calls for simple vs. tiered merging strategies for all autosomal DEL, DUP, and INV. For each merging strategy, bars indicate the number of SV overlapped (90% reciprocal overlap) by exactly by exactly 0, 1, 2-4, or >=5 variants called using the other strategy. (b) Comparison of CI length for simple versus tiered merged samples for each breakpoint of all autosomal DEL, DUP, and INV variants, restricted to variants (shown in red in panel a) with a 1-1 correspondence between merging strategies.

#### Supplementary Methods

##### Benchmarking sample selection

For both software and accuracy benchmarking of svtools, we used trios from four families, CEPH1463 (European), PR05 (Puerto Rican), SH032 (Han Chinese), Y117 (Yoruban) obtained from Coriell. In addition, we used 988 Finnish samples selected from the METSIM (Laakso, et al., 2017) (METabolic Syndrome In Men; 468 samples), FINRISK (Borodulin, et al., 2015) (478 samples), and EUFAM (Porkka, et al., 1997) (European Study of Familial Dyslipidemias; 42 samples) studies for benchmarking of larger cohorts. For the EUFAM samples, founders were selected. All samples were sequenced on the HiSeq X to a minimum depth of 20X with a contamination rate less than 5% as calculated by verifyBamId (Jun, et al., 2012) using population allele frequencies from the 1000 Genomes project (http://csg.sph.umich.edu/kang/verifyBamID/download/Omni25_genotypes_1525_samples_v2.b37.PASS.ALL.sites.vcf.gz). The three benchmarking cohorts were as follows: 10 samples (10 Coriell), 100 samples (12 Coriell + 88 METSIM) and 1000 samples (12 Coriell + 468 METSIM + 478 FINRISK + 42 EUFAM). For precision and sensitivity estimation, we used the 12 sample (12 Coriell) and 1000 sample (12 Coriell + 468 METSIM + 478 FINRISK + 42 EUFAM) cohorts.

##### Software performance benchmarking

In order to measure the compute resources necessary for each step of the svtools pipeline, we performed benchmarking using cohorts of 10, 100 and 1000 samples. Benchmarking was performed on a single, Ubuntu 10.04 node running within our IBM Platform LSF managed cluster. This node had two 4-core Intel Xeon E5420 @ 2.50Ghz processors, 32 GB of RAM, and a 146 GB, 10,000 RPM local hard drive with 16 MB of cache. Data and code (svtools 0.3.1) was first staged to the local disk to minimize the effect of network share performance on the benchmarks. Five replicates of each benchmark were performed serially and the first discarded, as it was intended to prime the filesystem cache. CPU, wall clock and memory metrics were gathered from IBM Platform RTM monitoring of individual benchmarking runs. Some benchmarking runs finished before LSF was able to gather memory usage metrics and these are reported as NA in Table 3.

##### Comparison to 1000 Genomes Project SV calls

We tested variant detection sensitivity utilizing calls from the 1000 Genomes Project (1KGP) Phase 3 integrated structural variation map (Sudmant, et al., 2015) (ftp://ftp.ncbi.nlm.nih.gov/1000genomes/ftp/phase3/integrated_sv_map/ALL.wgs.integrated_sv_map_v2.20130502.svs.genotypes.vcf.gz). To generate a “truth-set” of relevant variation that could potentially be detected using our tools, we extracted all SVs that were detected by read-pair and/or split-read alignment signals in 1KGP and that were from SV classes that our tools are designed to detect – namely, DELs, DUPs, INVs and reference genome MEIs. Since SV classification schemes vary between the studies, we performed comparisons based purely on the position and relative orientation of reported SV breakpoint coordinates and confidence intervals. To compare an individual sample's calls to its 1KGP calls, we first extracted variant sites for that sample from the 1KGP VCF and our own cohort VCF using bcftools (Li, 2011) (v1.3) and converted each to a sorted BEDPE file using svtools (v0.3.1). We determined whether a variant was shared between the two files using bedtools pairtopair (Quinlan and Hall, 2010) (v2.23.0), requiring that both breakpoint intervals from any two variant calls overlapped (allowing for 50 bp of "slop") with consistent breakpoint strand orientations. All code to both prepare the 1KGP calls and perform the comparisons is available at https://github.com/hall-lab/1kg_sv_comparison.

**Mendelian error rate**

Mendelian error (ME) rates were calculated by taking the number of mendelian errors reported by PLINK (Chang, et al., 2015; Purcell, et al., 2007) (v1.90b3.38) on chromosomes 1-22 (--mendel --allow-extra-chr --chr 1-22) divided by the number of informative variants in the family. Informative sites were calculated using bcftools (v1.3) as the number of variant sites in the trio where one or more parents had a homozygous genotype. In order to prevent double counting of BND variants, all BND lines marked as SECONDARY were removed before calculating ME rates.

**Generation of mobile element insertion annotation**

Mobile element insertion (MEI) annotations were generated from the UCSC Genome Browser's Repeatmasker table (Tyner, et al., 2017) (http://hgdownload.cse.ucsc.edu/goldenPath/hg19/database/rmsk.txt.gz). Briefly, we selected repeats, with less than 200 base mismatches per thousand bases, that were classified as SINEs, LINEs, or SINE-VNTR-Alu (SVA) retrotransposons. Code to generate the file is documented within the svtools tutorial (https://github.com/hall-lab/svtools/blob/master/Tutorial.md#generate-a-repeat-elements-bed-file).

**Generation of Build 37 exclude list**

To create a list of regions to exclude from structural variant calling, the reference was divided into 100 bp windows using bedtools makewindows and genotyped using cnvnator-multi (Abyzov, et al., 2011; Chiang, et al., 2015) across 3,015 samples from diverse ethnicities and sequencing projects. Exclusion regions were defined as those where the total copy number across all samples was greater than ten times the expected diploid copy number and accounting for expected deviations from diploidy due to sex.

**Generation of Build 38 exclude list**

For the GRCh38DH exclude list, we excluded any HLA, decoy or alternate contigs and regions of the genome where the total copynumber, as calculated by cnvnator (v0.3.3), across 409 diverse samples, was greater than 12 copies per genome, accounting for expected differences due to sex (https://github.com/hall-lab/speedseq/blob/master/annotations/exclude.cnvnator_100bp.GRCh38.20170403.bed).

**Generation of variant training set for variant classification**

The variants used to train the naïve Bayes classifier used in the small sample SV reclassification scheme was selected from a set of high-quality autosomal deletions and duplications in a previously-generated, post-quality control lumpy/svtools callset from 948 Finnish genomes. Variants with allele frequency >1%, and such that the mean copy numbers for heterozygotes and homozygous reference individuals fell between the 10$^{th}$ and 90$^{th}$ percentiles for all qc-passing autosomal variants of the same type, were selected from the Finnish callset. svtools varlookup was used to extract the same set of high-quality variants from the test data set, and the copy number profile for the set of overlapping variants was used for the svtools classify tool in naïve Bayes mode.

**Algorithm for variant reclassification**

Reclassification refines calls at copy number variants by incorporating read depth information at each putative DEL and DUP. Two algorithms are implemented: 'large sample' (intended for joint callsets of at least ~30 genomes) and 'naïve bayes' (appropriate for smaller joint callsets). The large sample algorithm examines each individual variant at a time, across all samples and regresses copy number against the estimated allele balance. If the direction of the slope is consistent with the SV class and the squared correlation between allele balance and copy number exceeds a user-defined value ($R^2$>0.2 by default), the classification of DEL or DUP is retained; otherwise, the variant class reverts to BND (i.e., a breakpoint of indeterminate type). For rare variants, read-depth support is assessed based on comparison of the copy number in each variant sample to the distribution of copy number values in wild-type samples.

The naïve bayes algorithm examines the copy number across all positions in the same sample. It is first trained based on the distribution of copy number values for each genotype class at a selected set of simple, high-quality copy number variants created as described above. Copy number values are assumed to be normally-distributed with mean dependent on the variant type and genotype, and pooled variance. Then for each putative CNV in the VCF, the variant either retains its class or is reclassified as a BND according to which maximizes the posterior probability in the naives bayes classifier.

During reclassification, variants with orientations indicative of deletions with a reciprocal overlap >0.9 to an annotated MEI element (as described above) were classified as MEIs (Chiang, et al., 2017).

**Generation of cohort-level VCFs used for assessing callset quality**

All samples were aligned to GRCh37-lite using speedseq (Chiang, et al., 2015) realign (commit 1aa63c99b02d76db58db1182efe450b27f98e819 at https://github.com/hall-lab/speedseq/tree/gms). For SV calling, individual samples were first processed through speedseq sv (v0.1.2) to generate structural variation calls with lumpy-sv (v0.2.11) followed by svtyper (v0.1.1) and cnvnator-multi histogram files. These per-sample files were then assembled into cohort-level VCFs using svtools (v0.3.1). Before merging, format fields were reordered using https://github.com/hall-lab/svtools/blob/develop/scripts/reorder_format_fields.py. VCFs were then sorted (svtools lsort) and merged with each call padded on either side by 20 bp (svtools lmerge -f 20 -g). Genotypes were preserved through merging to allow for quality evaluation.

After merging, each sample was genotyped using svtools genotype and copy number annotated using svtools copynumber. Per-sample VCFs were reassembled using svtools vcfpaste (-q -m lmerge.vcf). For the 1000 sample cohort, svtools reclassify -m large_sample was used to reclassify genotypes. For the 12 sample cohort, svtools reclassify -m naive_bayes was used to reclassify genotypes. To construct the training set, svtools varlookup was used to annotate 12 sample cohort calls overlapping a set of known, high quality variants (https://github.com/hall-lab/svtools/blob/develop/resources/training_vars.bedpe.gz) and the resulting intersected calls used as the training data.

After reclassification, variant calls were determined to be high-confidence based on the following filtering criteria: deletions, duplications, and mobile element insertions with mean sample quality ≥ 100; inversions with quality score ≥ 100, both evidence types (split-read or paired-end) providing ≥ 10% of the alternate allele's alignment support, and each respective breakpoint call providing ≥ 10% of the alternate allele's alignment support; breakends reclassified from deletion calls and with mean sample quality ≥ 500; or breakends not subject to reclassification with mean sample quality ≥ 250. All other variants were filtered out as low quality.

**Cost calculations**

Estimates of the per-sample cost of executing the GRCh38 pipeline on Google Cloud Platform were performed by using metadata available from the Google Genomics API to determine the instance type, persistent disk type, disk volume and runtime for a representative subset of data processed. These values were then used to estimate a total cost based on a Google Cloud Platform pricelist (v1.13 last updated on 01-August-2017) and divided by the number of samples in the workflow examined to get a per-sample cost. These estimates do not include network egress charges or sustained usage discounts. All code is available at https://github.com/ernfrid/cromwell_cost.
